# A SKI subcomplex specifically required for the degradation of ribosome-free RNA regions

**DOI:** 10.1101/409490

**Authors:** Elodie Zhang, Varun Khanna, Abdelkader Namane, Antonia Doyen, Estelle Dacheux, Bernard Turcotte, Alain Jacquier, Micheline Fromont-Racine

## Abstract

The Ski2-Ski3-Ski8 (SKI) complex assists the RNA exosome during the 3’-5’ degradation of cytoplasmic transcripts. Previous reports showed that the SKI complex is involved in the 3’-5’ degradation of mRNA, including 3’ untranslated regions (UTRs), devoid of ribosomes. Paradoxically, we recently showed that the SKI complex directly interacts with ribosomes during the co-translational mRNA decay and that this interaction is necessary for its RNA degradation promoting activity. Here, we characterized a new SKI-associated factor, Ska1, which antagonizes the SKI-ribosome interaction. We showed that the SKI-Ska1-subcomplex is specifically involved in the degradation of ribosome-free RNA regions such as long mRNA 3’UTRs and cytoplasmic lncRNAs. We propose a model in which the SKI-exosome complex first targets ribosome-free RNA 3’ends in its Ska1-associated form. When the complex reaches the mRNA coding sequence, the Ska1-SKI-exosome complex is exchanged for the SKI-exosome, which interacts directly with ribosomes in order to resume the degradation process.

## Introduction

RNA degradation is a key component of RNA homeostasis and quality control. During normal mRNA turnover, controlled deadenylation is followed by 5’-3’ mRNA degradation by the cytoplasmic exonuclease Xrn1 as well as 3’-5’ degradation involving the cytoplasmic exosome (see for review Fromont-Racine and Saveanu, 2014; Parker, 2012). In the major 5’-3’ mRNA degradation pathway, mRNA deadenylation is followed by decapping by Dcp1/Dcp2 with the help of several enhancers of decapping such as Edc1, Edc2, Edc3, Dhh1, Pat1, Scd6 and the Lsm complex. This is followed by mRNA degradation *via* Xrn1, a co-translational process (Hu *et al*, 2009; Pelechano *et al*, 2015). The 3’-5’ mRNAs degradation pathway involves the cytoplasmic exosome composed of ten main factors, including the catalytically active subunit Dis3 (Allmang *et al*, 1999). Depending on its localization, the exosome is assisted by different co-factors. The nuclear exosome associates with the distributive exoribonuclease Rrp6 as well as Mpp6, Lrp1/Rrp47 and the RNA helicase Mtr4, which is found within the TRAMP complex (Trf4/5, Air1/2, Mtr4) (de la Cruz *et al*, 1998; Mitchell *et al*, 2003; Milligan *et al*, 2008). In the cytoplasm, the exosome is targeted to specific substrates by the SKI complex, composed of Ski2, Ski3 and Ski8 (SKIV2L, TTC37 and WDR61, respectively, in human).

The *SKI* genes were identified 40 years ago by performing a genetic screen with a yeast strain containing linear double-stranded RNAs encapsulated into virus-like particles secreting toxic proteins. Mutants having a deleterious phenotype were isolated and called *SKI* for super killer (Toh-E *et al*, 1978). Therefore, the first function reported for the *SKI* genes was in antiviral defense through viral RNA degradation (Widner & Wickner, 1993). Twenty years later, these genes were shown to be required for mRNA 3’-5’ degradation (Anderson & Parker, 1998) and to form a complex *in vivo* (Brown *et al*, 2000). Native mass spectrometry analyses revealed that the yeast SKI complex is a heterotetramer formed of one copy of Ski2 and Ski3 and two copies of Ski8 (Synowsky & Heck, 2008). Ski2 is an RNA helicase with a structure similar to the nuclear Mtr4 helicase. Ski8 contains WD40 repeats while Ski3 is a tetratricopeptide repeat (TPR)-containing protein, which forms the scaffold of the SKI complex (Halbach *et al*, 2013). Ski2 is thought to be required to unwind RNA and to remove proteins bound to mRNAs. Ski3 and Ski8 contribute to the structure and activity of the SKI complex (Halbach *et al*, 2013). Although Ski7 is not part of the SKI complex, it is also involved in mRNA 3’-5’ degradation (van Hoof *et al*, 2000) through bridging the SKI complex and the exosome (Araki *et al*, 2001). Interestingly, Ski7 interacts with the same exosome region as the nuclear factor Rrp6 (Kowalinski *et al*, 2016). Recently, structural data showed that the SKI complex forms a path for the RNA through a central channel of its helicase, Ski2. RNase protection assays performed with a mixture of the exosome and the SKI complex revealed protected fragments with a length compatible with a continuous channel formed by the SKI and the exosome complexes. It was proposed that the SKI complex directly delivers single-stranded RNA into the exosome (Halbach *et al*, 2013). Altogether, these results led to a model in which the cytoplasmic exosome and the SKI complex could be structurally organized to degrade RNA molecules, in a fashion similar to the proteasome involved in protein degradation (Halbach *et al*, 2013).

In addition to its role in the general mRNA degradation pathway, the SKI complex is also required for cytoplasmic mRNAs surveillance pathways from yeast to humans: Non-Stop mRNA decay (NSD), No-Go mRNA decay (NGD) and Non-sense mediated mRNA decay (NMD) (Frischmeyer, 2002; van Hoof *et al*, 2002; Mitchell & Tollervey, 2003; Doma & Parker, 2006). Recent *in vivo* and *in vitro* experiments have shown that the 3’-5’ mRNA degradation of mRNAs by the SKI-exosome complex is a co-translational event (Schmidt *et al*, 2016), in analogy to the Xrn1 dependent 5’-3’ mRNA degradation pathway (Pelechano *et al*, 2015). This conclusion is supported by a series of observations: *i*) affinity purification of a nascent peptide captured on a ribosome stalled at the end of a non-stop mRNA co-purifies the ribosome in association with the SKI complex; *ii*) a Ski2 affinity purification enriches the ribosome together with the SKI complex; *iii*) these SKI-ribosome complexes are found associated with most open reading frames, as determined by footprinting experiments; *iv*) the cryo-electron microscopy structure of the SKI-ribosome complex shows that the SKI complex interacts intimately with the small subunit of the ribosome. Notably, this interaction appears to be important to activate the Ski2 helicase by changing the conformation of two elements that autoinhibit the ATPase activity of the SKI complex. The arch domain of Ski2 moves away from the RNA helicase channel and the N-terminal arm of Ski3 adopts a more open conformation (Schmidt *et al*, 2016). However, these observations are at odd with previous reports which showed that the SKI complex is required for the efficient degradation of 3’UTRs by the exosome (Anderson & Parker, 1998). Since 3’UTRs are essentially devoid of ribosomes, this implies that the SKI complex can act as an exosome co-factor without requiring an association with the ribosomes. In an attempt to solve this paradox, we investigated the mechanism of action of the SKI complex on ribosome-free RNA regions.

Here, we have identified a new component of the SKI complex that we called Ska1 for SKI-Associated component 1. This protein is encoded by *YKL023W*, a gene of unknown function. We demonstrated that this factor is not involved in the translation-dependent NSD pathway but rather in the general RNA degradation pathway, specifically in the degradation of RNA regions devoid of ribosomes. The SKI-Ska1 subcomplex is required for the 3’-5’ degradation of mRNA 3’UTR regions and cytoplasmic long non-coding RNAs (lncRNAs). Interestingly, Ska1 is dispensable for the efficient degradation of coding mRNA regions or for the elimination of aberrant transcripts such as non-stop mRNAs. Overexpression of *SKA1* antagonizes the association of the SKI complex with ribosomes, precluding its activity in degrading ribosome associated RNA regions such as NSD substrates. These results show that, depending on the translation status of the RNA region being targeted, the SKI complex interacts either with the ribosome or Ska1. Association of Ska1 with SKI promotes loading on ribosome-free RNA 3’ ends and stimulates degradation by the exosome.

## Results

### Ska1 is a novel factor associated with a subpopulation of the SKI complex

To investigate the mechanism of action of the SKI complex, we performed an affinity purification using Ski3-TAP as a bait. The proteins associated with Ski3-TAP were separated on a polyacrylamide gel (Fig 1A) and identified by mass spectrometry (LC-MS/MS). Analysis of the label-free quantitative MS data, represented by a volcano-plot, showed the enrichment for three groups of proteins associated with Ski3 (Fig 1B). As expected, the most enriched proteins relative to the untagged strain control were the bait Ski3-TAP and the two other subunits of the SKI complex, Ski2 and Ski8. A second group was composed of the six RNasePH domain proteins of the exosome (Rrp41, Rrp42, Rrp43, Rrp45, Rrp46, Mtr3, and the catalytic subunit Dis3) and of its three RNA-binding subunits (Rrp4, Rrp40 and Csl4). These factors form the 10-subunit cytoplasmic exosome (Fig. 1B, in yellow) (Liu *et al*, 2006). Ski7 and a novel factor of unknown function, Ykl023w, were also enriched at a similar level than the exosome subunits. We named the uncharacterized Ykl023w protein Ska1 for SKI-Associated component 1 (Fig 1B). The third group of proteins was mainly related to the ribosome.

**Figure 1.**
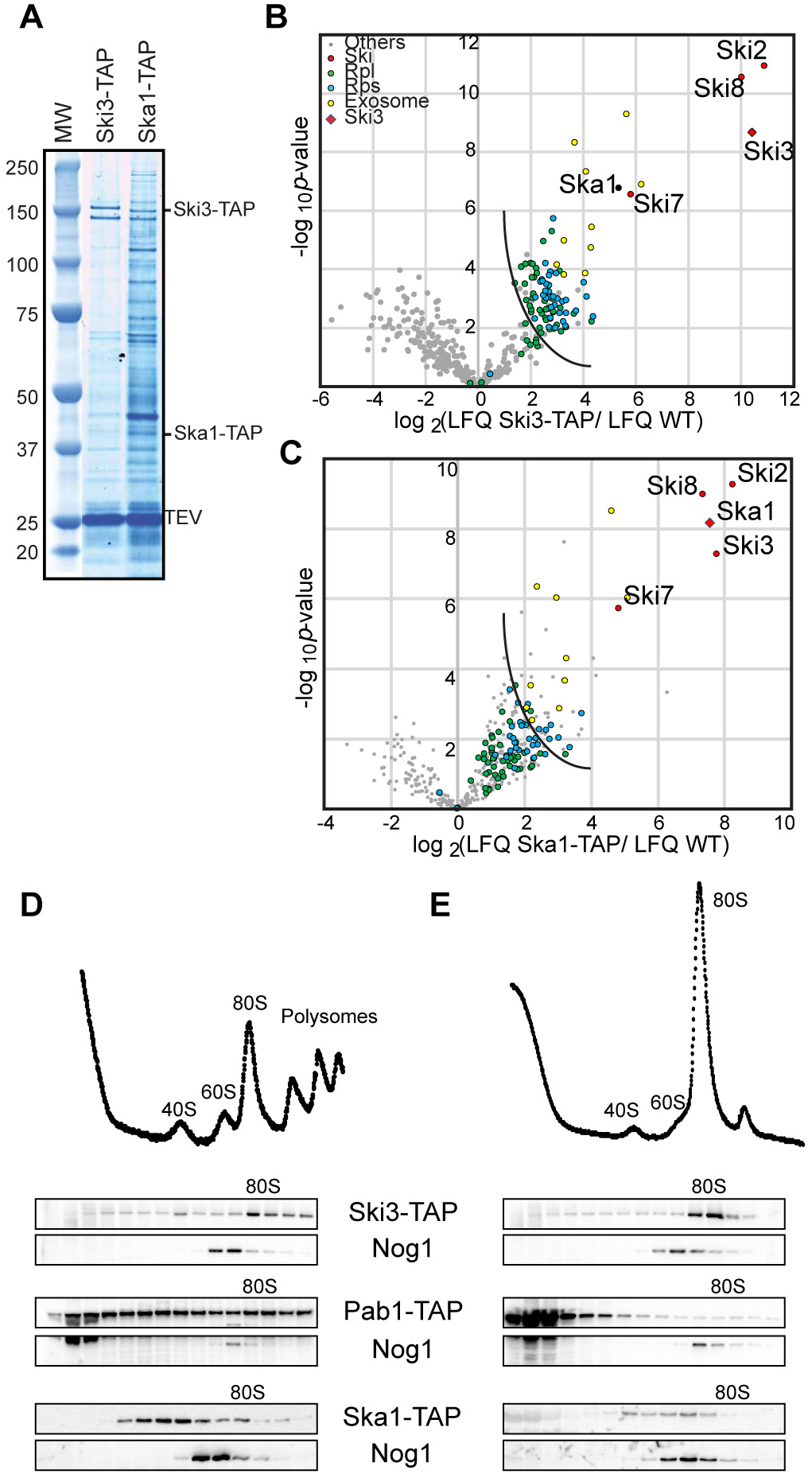
Ska1 is associated with the SKI complex. **A.** Affinity purification using Ska1-TAP and Ski3-TAP. The components of the TEV eluates were separated on a polyacrylamide NuPAGE Novex 4-12% Bis-Tris gel (Life Technologies) and visualized with Coomassi blue staining. MW: Molecular Weight marker. **B.** Statistical analysis of the fold enrichment of protein after one-step affinity purification using the Ski3-TAP as a bait (six replicates) followed by mass spectrometry analysis (LC-MS/MS). The enrichment was calculated relative to a control purification using the untagged reference strain BY4741 and represented, on the x-axis, as log2(LFQ Ski3-TAP/LFQ WT) where LFQ stands for Label Free Quantification (LFQ; Appendix). The y-axis represents the *p* value distribution (-log10 *p* value) calculated using the Student’s t test for all identified proteins represented by a circle. Red; the SKI complex; yellow, the exosome; blue, the small subunit ribosomal proteins - RPSs; green, the large subunit ribosomal proteins - RPLs). The diamond indicates the protein used as a bait for the affinity purification. Proteins above the curved lines show a statistically significant enrichment according to the t-test value (see dataset EV1). **C.** Same as in (B) but for the Ska1-TAP purification (six replicates). **D.** Total cellular extracts from cells expressing either Ski3-TAP or Ska1-TAP or Pab1-TAP were separated on a sucrose gradient (10-50%) by ultracentrifugation. Proteins of each fraction were analyzed by Western blot using a PAP antibody for the detection of the TAP fusion protein and anti-Nog1 antibody to mark the 60S fraction. **E.** Total cellular extracts were treated with RNase before loading on the sucrose gradient and analyzed as in (D).

To confirm the physical interaction between Ska1 and the SKI complex, we performed affinity purification using a Ska1-TAP strain in the same conditions of purification used for Ski3-TAP (Fig 1C). We tested the functionality of the TAP fusions combining Ski3-TAP or Ska1-TAP with a Dcp2-degron mutant (see below). We observed that Ska1-TAP was partially functional in this assay in contrast to the fully functional Ski3-TAP (Fig EV1). Nevertheless, the analysis of label-free quantitative MS data revealed that the complex associated with Ska1 was highly similar to the SKI complex purification. We again observed an enrichment for three groups of proteins. The most enriched proteins were Ski2, Ski3, and Ski8, which were enriched to levels comparable to the Ska1-TAP bait. As observed with Ski3-TAP, the second group of proteins associated with Ska1-TAP was mainly related to the exosome and Ski7. The enrichment for the third group of proteins related to the ribosome was less important than the one observed with the Ski3-TAP purification.

To better evaluate the relative distribution of Ska1 and SKI components in the analyzed complexes, we looked at the relative protein abundance in a wild-type yeast strain (Nagaraj *et al*, 2012). The comparison of the abundance of 4077 proteins revealed that Ska1 and SKI complex were expressed at low levels. In addition, Ska1 was expressed to a lower level compared to the components of the SKI and exosome complexes (Fig EV2 upper panel). However, despite its lower abundance, Ska1 was enriched to a level comparable to the exosome components after purification of the complex associated with Ski3 (Fig EV2 middle panel). In addition, the Ski proteins were more enriched in the Ska1-TAP than the exosome (Fig EV2 lower panel). These results suggest that most of the Ska1 molecules are associated with the SKI complex. In contrast, only a fraction of the SKI complex was associated with Ska1.

Affinity purification showed that Ska1 was also associated with the ribosome, yet to a lesser extent compared to Ski3 (Fig 1B and 1C). To verify this association, we fractionated cellular extracts from a Ska1-TAP strain on a sucrose gradient. The analysis of the fractions by Western blot revealed that Ska1-TAP co-sedimented with the 80S peak, the light polysomal fractions and lighter fractions, in a region of the gradient where the 40S and 60S subunits sediment (Fig 1D). For comparison, a similar experiment, performed with the Ski3-TAP strain showed a greater enrichment of Ski3 in the 80S peak, as previously described (Fig 1D) (Schmidt *et al*, 2016). These observations are in agreement with the results of affinity purification showing a stronger association of Ski3-TAP than Ska1-TAP with the ribosome. In order to determine if the weak association of Ska1 with the 80S ribosome and polysomes was direct or mRNA-mediated, we performed a RNase treatment of Ska1-TAP, Ski3-TAP cellular extracts before their fractionation on a sucrose gradient (Fig 1E). In addition, we used Pab1-TAP as a mRNA-binding protein control. As expected, RNase treatment resulted in the disappearance of the polysomes and the accumulation of 80S monosomes. After RNAse treatment, Ski3-TAP was still associated with the 80S ribosome, as previously shown (Schmidt *et al*, 2016), whereas Pab1-TAP, which was distributed all along the sucrose gradient in absence of RNase, sedimented in the light fractions as a free protein, thus confirming the efficiency of the RNase treatment. In contrast, Ska1-TAP showed a sedimentation profile different from both Ski3-TAP and Pab1-TAP profiles. We observed an enrichment for Ska1 in the fractions mainly sedimenting around the 40S and 60S peaks but not within the 80S peak (Fig 1E).

These results indicated that following RNase treatment, Ska1 was still associated with large complexes but not with the 80S ribosome and polysomes, suggesting that the weak association of Ska1-TAP to the ribosome observed in the affinity purification was RNA-dependent. The presence of Ska1-TAP in several fractions of the gradient suggested that Ska1 could be associated with different complexes. Altogether, these results suggest that there are at least two different SKI complexes with one containing Ska1.

### Ska1 is involved in mRNA 3’UTR degradation but not NSD

Since Ska1 physically interacts with the SKI complex, we determined if Ska1 participates in the same pathways as the SKI complex. The SKI complex contributes to the degradation of a number of RNA substrates, including conform mRNAs but also aberrant ones such as non-stop mRNAs. To investigate if Ska1 is also involved in NSD, we tested the expression of the TAP-Non-Stop and GFP-Rz-HIS3 reporter genes (Defenouillère *et al*, 2013; Kobayashi *et al*, 2010) in the presence or the absence of Ska1 (Fig 2A left and right panels). Both reporters are sensitive to the NSD pathway. The TAP-Non-Stop reporter gene codes for Protein A, but lacks a stop codon. The polyA tail is thus translated into a poly–lysine stretch, which, due to its high electrostatic charge, gets stuck at the ribosome exit tunnel, resulting in translation stalling. In contrast, the GFP-Rz-HIS3 reporter gene carries a ribozyme, which generates an mRNA coding for GFP without a stop codon and a polyA tail. As a result, translation stops only when the ribosome reaches the very end of the transcript. We performed Western blot analyses with total cellular extracts from the *ska1*Δ or the *ski3*Δ mutants and wild-type strains transformed with plasmids expressing these reporters. As controls, we used the same reporter genes but with a normal termination codon or without a ribozyme. We did not observe any aberrant protein accumulation in the *ska1*Δ strain. In contrast, the *ski3*Δ strain positive control exhibited a strong accumulation of these aberrant polypeptides (Fig 2A). These results indicated that Ska1 is not involved in the Non-Stop aberrant mRNA degradation pathway.

**Figure 2.**
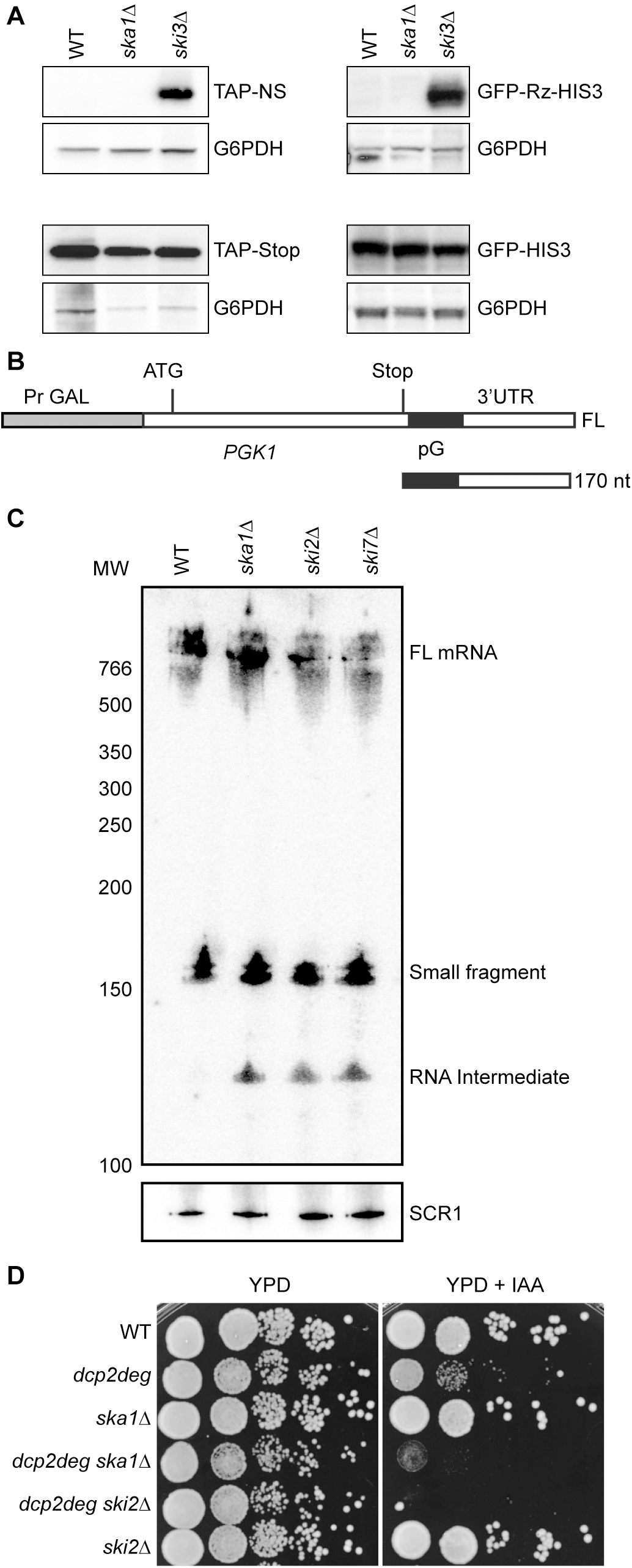
Ska1 is involved in the 3’-5’ mRNA degradation pathway. **A.** Ska1 is not required for Non-Stop mRNA degradation. Total cellular extracts were prepared from strains transformed with the reporter plasmids TAP-Non-Stop, GFP-Rz-HIS3, TAP-Stop or GFP-HIS were separated on a 10% SDS-PAGE polyacrylamide gel. Protein A and GFP were revealed by Western blot using a PAP antibody and an anti-GFP antibody, respectively. A loading control was obtained using an anti-G6PDH antibody. **B.** Schematic representation of the *PGK1* reporter gene (Anderson & Parker, 1998). Top: the grey box represents the *GAL1* promoter region (PrGAL), the white boxes the *PGK1* transcribed sequences and the black box the polyG tract inserted just downstream of the *PGK1* stop codon. Bottom: the 170 nucleotides fragment that accumulates as a result of the blockage of the 5’-3’ exonucleolytic degradation by the polyG sequence. **C.** Total RNA from wild-type, *ska1Δ, ski2*Δ and *ski7*Δ strains was separated on a 6%-Urea polyacrylamide gel. Northern blot was performed with a [^32^P]-polyG probe. The membrane was then hybridized with a [^32^P]-radiolabeled *SCR1* oligonucleotide as a loading control. MW: positions of the DNA fragments (in nucleotides) used as molecular weight ladder. **D.** *ska1*Δ is synthetic lethal with the *dcp2degron* mutant. Serial dilutions of yeast strains, as indicated, were spotted on rich medium plates in the presence or absence of auxin (0.1 mM) and incubated for 48h at 30°C.

To test the effect of *SKA1* removal on the general mRNA degradation pathway, we followed the behavior of the well-characterized reporter *PGK1pG* expressing a stable transcript (Anderson & Parker, 1998). This gene is under the control of the *GAL1* promoter, which is active in the presence of galactose but totally repressed with glucose. Addition of glucose allows to monitor the rate of degradation of the reporter mRNA. The stop codon is followed by a polyG tract which blocks the progression of Xrn1 in the 3’UTR allowing the detection of 3’-5’ degradation intermediates (Fig 2B) (Anderson & Parker, 1998). With this *PGK1pG* reporter gene, the full-length mRNA is detected as well as a small 3’UTR fragment that accumulates as a result of the polyG blocking Xrn1. As previously observed (Anderson & Parker, 1998), the absence of Ski2 or Ski7 resulted in the accumulation of a short 3’UTR degradation intermediate, indicative of the inefficient 3’-5’ degradation in the absence of a functional SKI complex. Importantly, the absence of Ska1 resulted in the accumulation of the same 3’UTR intermediate fragment (Fig 2C). This result indicates that Ska1 is involved in the same general mRNA decay mechanism as the Ski factors concerning the 3’UTR regions.

If Ska1 is involved in the 3’-5’ degradation mRNA pathway, we postulated that the absence of Ska1 combined with mutation into genes involved in the 5’-3’ mRNA degradation pathway should be deleterious for cells. Since the absence of the decapping enzyme Dcp2 severely affects cell growth, we fused a degron sequence to *DCP2* to induce its degradation in the presence of auxin (IAA). We combined this *DCP2* mutant version with a *ski2*Δ or *ska1*Δ mutation. Growth of *ska1*Δ or *ski2*Δ mutant strains was not affected as compared to a *dcp2degron* strain in the presence of auxin. In contrast, the double-mutant strains were sensitive to auxin (Fig 2D). We observed a synthetic lethality phenotype for cells combining *ski2*Δ and the *dcp2degron* mutant, as previously described (Anderson & Parker, 1998). Although less severe, a *ska1*Δ and *dcp2degron* strain also showed a marked growth defect.

### Ska1 is specifically required for the 3’-5’ degradation of ribosome-free RNA regions

Since we observed intermediates of degradation accumulating in the absence of Ska1 or Ski2 using a *PGK1pG* reporter gene (Fig 2C), we analyzed the genome-wide effect of *SKA1* removal. To detect any deficiency of the 3’-5’ degradation pathway, it is necessary to inhibit the major 5’-3’ pathway. Upon *XRN1* deletion, mRNAs accumulate as decapped molecules, while the deletion of *DCP2* or *DCP1* severely impairs growth. In order to preserve more physiological conditions at the time of analysis, we used a *dcp2degron* mutant which, upon auxin addition, strongly decreases mRNA decapping, protecting mRNAs 5’-ends from the Xrn1 exonucleolytic degradation (see Fig 2D and Fig EV3A). We mapped and quantified RNA 3’-ends genome-wide in a wild-type and in single and double mutant strains combining the *dcp2degron* mutant with *ska1*Δ or *ski2*Δ mutations. We used an anchored oligo-dT cDNA-priming step to generate 3’-end RNA sequencing libraries (see Materials and Methods). Since the 3’-5’ RNA degradation intermediates are predicted to be devoid of polyA, we added an *in vitro* polyadenylation step using polyA polymerase. RNAs were extracted after two hours of auxin treatment. Note that the vast majority of the Dcp2 protein was already depleted after one hour of auxin treatment (Fig EV3A). However, we performed RNA analysis after two hours of treatment in order to allow degradation intermediates to accumulate (see below). Examining the results with the Integrative Genomics Viewer (IGV) (Robinson *et al*, 2011), one of the most striking phenotype observed with the *dcp2degron ski2*Δ double mutant in comparison with the *dcp2degron* single mutant was an accumulation, for many mRNAs, of trimmed 3’ ends over a distance of about 50 nucleotides from the polyA sites (Fig 3A and 3B and Fig EV3B). These 3’ trimmed transcripts were not visible in the *dcp2degron* or *ski2*Δ single mutants. We interpreted these shortened transcripts as 3’-5’ RNA degradation intermediates that accumulated over the Dcp2 depletion period (2 h in this experiment) when the 3’-5’ exonucleolytic activity of the exosome is very inefficient as a result of the SKI complex disruption in the *ski2*Δ mutant. Importantly, similar degradation intermediates were also observed in the *dcp2degron ska1*Δ double mutant, confirming on a genome-wide scale that Ska1 is important for the 3’-5’ mRNA decay pathway, in agreement with the observation made with the *PGK1pG* reporter gene (Fig 2C). Although the deletion of either *SKA1* or *SKI2* in a *dcp2degron* background showed similar effects on relatively long 3’UTRs (Fig 3A), removal of Ska1 did not result in changes for most mRNAs with very short (less than ∼60-70 nucleotides) 3’UTRs (Fig 3B and Fig EV3B).

**Figure 3.**
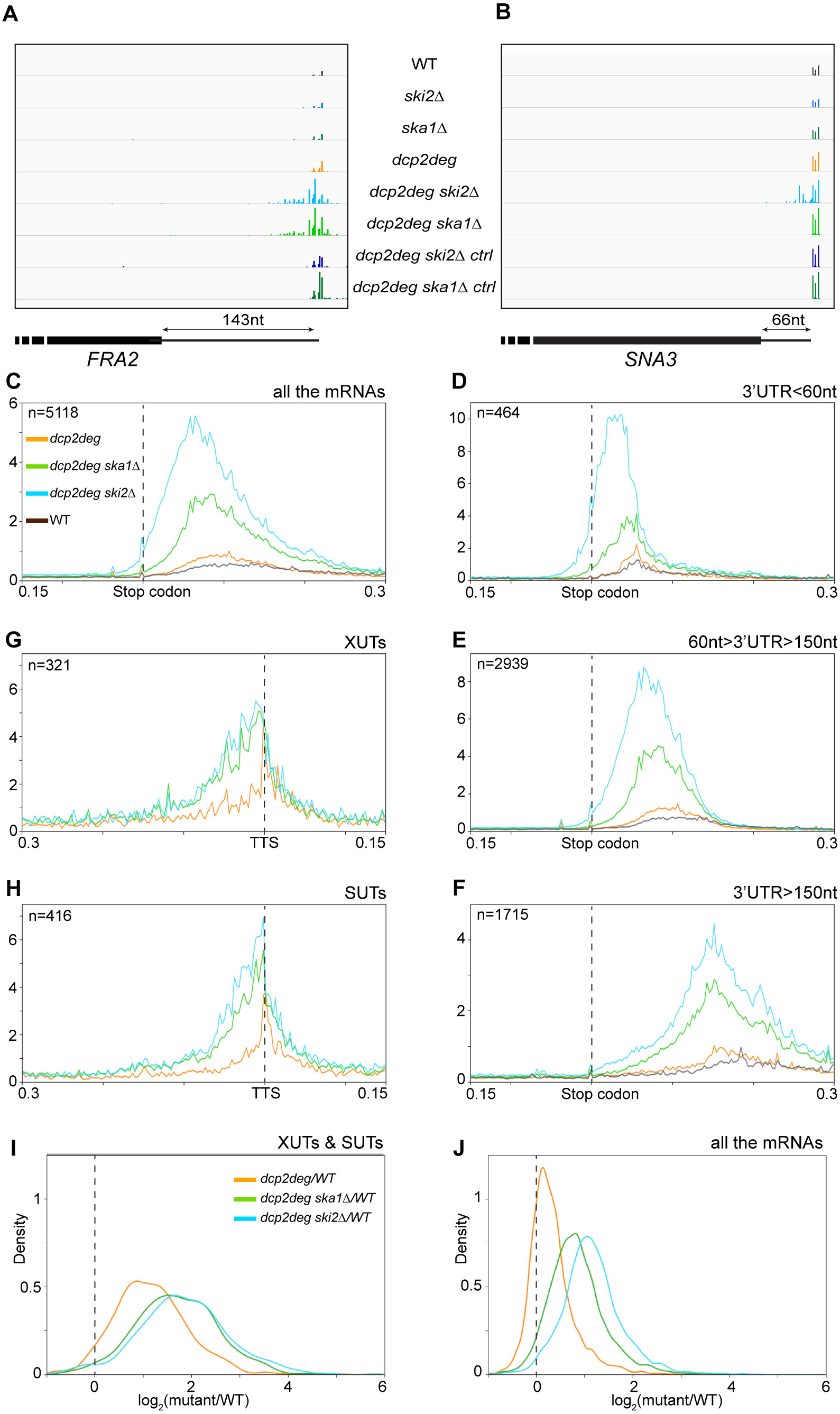
Ska1 and the SKI complex are required to degrade cytoplasmic RNA sequences devoid of ribosomes. **A-B.** Visualization of two loci, *FRA2* (long 3’UTR) and *SNA3* (short 3’UTR), using IGV to show the accumulation of 3’ end mRNA intermediates in the absence of the SKI complex or in the absence of Ska1 when the 5’ end degradation pathway is compromised. The name of the strains is indicated for each row, with “ctrl” standing for “control experiment without the *in vitro* polyadenylation step”, allowing the visualization of the naturally polyadenylated 3’ ends. The scheme at the bottom of the figure depicts the 3’ part of the genes with the thick box representing the end of the ORF and the line the 3’UTR. **C.** Genome-wide analysis of 3’-end mRNAs with protein-coding genes aligned with their stop codons. The curves indicate the distribution of 5’-end read counts for each condition, computed for each position ranging from -150 to +300 nucleotides relative to the stop codons (see Materials and Methods for more details). The distributions correspond to the average distributions of 3 independent replicates (see Fig EV5B for the data from all single experiments). **D-F.** As in (C) but for three groups of genes classified according to their 3’UTR length. **G-H.** As in (C) but for XUTs and SUTs aligned on the transcription termination sites (TTSs) and for positions -300 to +150 nucleotides relative to the TTS. **I-J.** Frequency distributions of the ratios of RNAseq read counts for mutant mRNAs compared with wild-type for mRNAs (J) and XUTs and SUTs (I), respectively.

Using Northern blot analysis upon RNase H digestion, we confirmed that the extent of 3’ end trimming was dependent on the duration of Dcp2 depletion (Fig EV3C; note that the *RPS31* gene belongs to the “short 3’UTR” category, not sensitive to Ska1). Thus, these shortened transcripts most likely correspond to 3’-5’ RNA degradation intermediates that accumulated following Dcp2 depletion.

We investigated the correlation between the length of the 3’UTR and the effect of Ska1 removal on the 3’-5’ degradation. To this end, we performed genome-wide analysis on total mRNAs and on mRNAs divided into three groups according to their 3’UTR lengths: genes with short 3’UTRs (16 to 60 nucleotides), genes with 3’UTRs of intermediate lengths (60 to 150 nucleotides), and genes with 3’UTRs longer than 150 nucleotides (Fig 3C-F). In order to specifically analyze the degradation intermediates, we eliminated from the analyses the normal cleavage and polyadenylation sites (i.e. the *bona fide* cleavage and polyA sites) as determined with the samples in which the *in vitro* polyadenylation step was omitted. Furthermore, to avoid the metagene signal being dominated by a few strongly expressed genes, the number of reads were normalized by the relative expression of the corresponding genes, as determined by standard RNAseq (see Materials and Methods). Fig 3F shows that long 3’UTRs shared a similar profile of accumulation of 3’ trimmed RNA intermediates in the absence of Ska1 or Ski2. In contrast, in the group of short 3’UTRs (less than 60 nucleotides) (Fig 3D), the degradation intermediates accumulated to a much lower extent in the absence of Ska1 when compared to the absence of Ski2. The group of genes with intermediate 3’UTR lengths exhibited an intermediate phenotype (Fig 3E). Note that Ska1 still retained some activity for mRNAs within the “short 3’UTR” class (Fig 3D). One possible explanation for this observation is that some mRNAs are poorly translated, thus requiring Ska1 for their 3’-5’ degradation by the exosome. For example, uORFs containing mRNAs are known to be generally poorly translated (Arribere & Gilbert, 2013). Figure EV4A shows three uORFs containing mRNAs with short 3’UTRs. These mRNAs behave as XUTs and SUTs and Ska1 is therefore as important as Ski2 for their efficient 3’-5’ degradation. However, normally translated mRNAs with 3’UTRs shorter than 60 nucleotides generally do not require Ska1 for their 3’-5’ degradation by the SKI-exosome complex. The fact that Ska1 was not required for efficient 3’-5’ degradation of short 3’UTRs was consistent with the observation that a NSD reporter construct, which has no 3’UTR, was not sensitive to the absence of Ska1, in contrast to the *PGK1pG* reporter transcript (Fig 2A). Naturally occurring premature polyadenylation sites, located within coding sequences, have been described for genes such as *RNA14* and *AEP2* (Sparks & Dieckmann, 1998). Such prematurely terminated transcripts have all the attributes of natural NSD substrates. Indeed, we observed a strong accumulation of the corresponding transcripts in the absence of Ski2 but not in the absence of Ska1, both within the 3’end RNAseq data, visualized using IGV, and Northern blots (Fig 4A-D). Furthermore, these accumulated transcripts were already detected in the 3’end RNAseq data without *in vitro* polyadenylation, demonstrating that they were polyadenylated *in vivo* (Fig 4A-D). These transcripts accumulated upon *SKI2* deletion to a much greater extent in a *dcp2degron* background than in the wild-type background, showing that the 5’-3’ degradation pathway remains important even for these non-stop transcripts.

**Figure 4.**
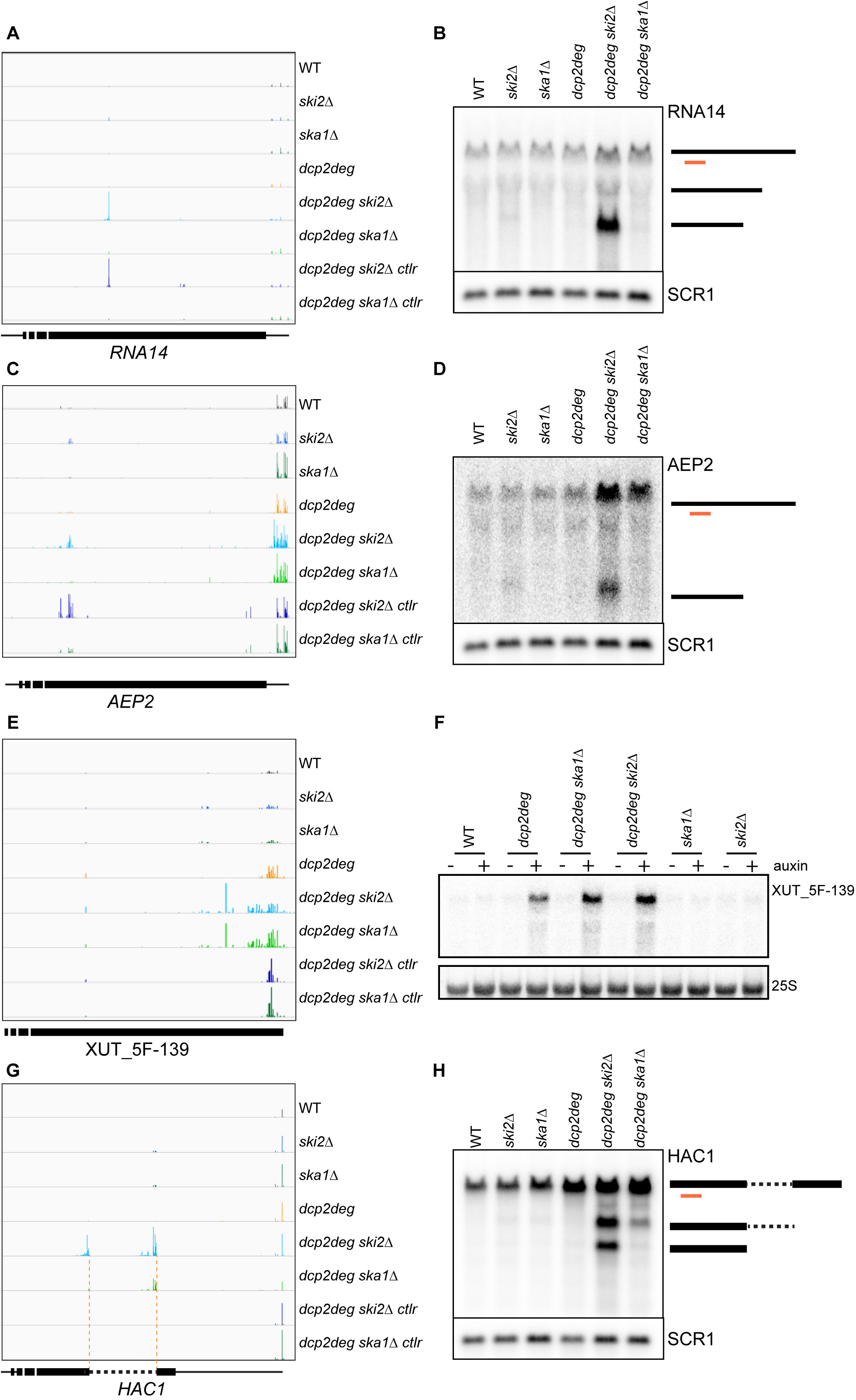
Ska1 is required for the degradation of XUTs but not of NSD targets. **A.** Visualization of the *RNA14* locus using IGV. Rows labeled as in Fig 3. **B.** Northern blot of total RNAs from strains treated with auxin for 2h. RNAs were separated, after denaturation at 65°C in formazol, on a 1% agarose gel and the blot hybridized with a [^32^P]-radiolabeled *RNA14* RNA probe (see Appendix) and then with a [^32^P]-radiolabeled *SCR1* oligonucleotide probe as a loading control. The different forms of the mRNA are schematized on the right by black lines, with the position of the RNA probe in orange. **C-D.** as in (A-B) but for the *AEP2* transcript. **E-F.** as in (A-B) but for the *XUT_5F-139* transcript; the Northern blot was performed with total RNAs from yeast strains treated or not with auxin for 2h. An rRNA 25S oligonucleotide probe was used as a loading control. **G.** Visualization of the *HAC1* locus in the different strains using IGV. The orange dotted lines indicate the splicing sites. **H.** as in (B) but for the *HAC1* transcripts. The dotted and thick black lines on the right schematize the intron and exon sequences, respectively.

We have previously shown that NSD substrates are associated with the ribosome (Schmidt *et al*, 2016). These results suggested that Ska1 could be specifically required with the SKI complex to target cytoplasmic RNA regions devoid of ribosomes. To analyze a set of transcripts with little association with ribosomes, we examined lncRNAs such as the XUTs and the SUTs, which are cytoplasmic pervasive transcripts containing only short spurious ORFs and very long untranslated 3’ regions. We aligned the reads of the XUTs or SUTs on their transcription termination sites (TTSs). We observed a similar profile of 3’ trimming in the absence of Ska1 and in the absence of Ski2, which was fully consistent with our hypothesis (Fig 3G and 3H). The observation that the deletion of *SKA1* or *SKI2* had similar effects on the accumulation of XUTs or SUTs was confirmed by an RNASeq approach (Fig 3I). We used the TruSeq stranded mRNA sequencing kit to construct RNA libraries and quantified transcripts in various strains. Since the mutants of interest are involved in the classical mRNA degradation pathway, we added *S. pombe* cells to the *S. cerevisiae* cell pellet for spiked-in normalization. We impaired the 5’-3’ pathway by depleting Dcp2 in strains lacking *SKA1* or *SKI2*. We clearly observed a similar accumulation of the XUTs and SUTs in these strains. In contrast, mRNAs accumulated more in the absence of Ski2 than in the absence of Ska1 (Fig 3J). This observation is consistent with the synthetic lethal effect on the growth that we observed previously (Fig 2D). Using IGV, we focused on specific XUTs and verified their enrichment by Northern blot (Fig 4E and 4F). These transcripts accumulated upon depletion of Dcp2 and the effect was more marked when the depletion of Dcp2 was combined with a deletion of *SKI2* or *SKA1*.

The *HAC1* gene is of particular interest to discriminate between the effect of Ska1 on ribosome associated and ribosome-free RNA 3’ends. The *HAC1* transcript contains a peculiar intron, which is cleaved by a splicing endonuclease complex generating two cleavages, one at the end of exon 1 and one at the end of the intron (Sidrauski et al., 1996). The cleaved 3’end of exon 1 is associated with a ribosome but not the intron (Guydosh & Green, 2014). We observed that the removal of Ska1 resulted in the accumulation of an exon 1-intron intermediate while no effect on exon 1 was observed. In contrast, the absence of Ski2 impaired the degradation of both transcripts (Fig 4G and 4H). Altogether, these results show that Ska1 is required for the efficient 3’-5’ degradation of ribosome-free RNA regions but not for the degradation of RNA regions associated with ribosomes, such as NSD substrates or very close to their 3’end (mRNAs with short 3’UTRs) (Fig EV3B).

The different effect of Ska1 relative to the 3’UTR length could be explained by the presence of two distinct SKI associated complexes. Indeed, affinity purification experiments led to the identification of a SKI complex associated with the ribosome and one associated with Ska1. The fact that very short 3’UTRs were not dependent upon the presence of Ska1 for their degradation suggested that, in contrast to longer 3’UTR, the presence of the ribosomes in close proximity with the 3’ end of the transcripts allows the formation of an active ribosome/SKI/exosome complex (Schmidt *et al*, 2016) by recruitment of a SKI complex to the terminal ribosome on the translated ORF. In contrast, when the last ribosome is away from the mRNA 3’end, as is the case with mRNAs with long 3’UTRs, Ska1 is required for the SKI complex to target the RNA 3’ends in the absence of a direct interaction with the terminal ribosome.

### Two different SKI complexes for two different classes of substrates

All our results point to the existence of two distinct SKI complexes: one containing Ska1 to specifically target ribosome-free RNA 3’ends and one lacking Ska1, which allows its direct association with the ribosome. In this model, Ska1 may act as a specific competitor for binding to the SKI ribosome-interacting surface, preserving a fraction of the SKI complex from interacting with ribosomes and allowing it to directly target ribosome-free RNA 3’ends. According to this model, overexpression of Ska1 should prevent SKI complexes from interacting with ribosomes, which should specifically affect the NSD degradation pathway but not the degradation of ribosome-free RNA regions.

To test this prediction, specifically the potential competition between Ska1 and ribosomes for the activation of the SKI complex, we investigated the effects of *SKA1* overexpression on RNA degradation. *SKA1* overexpression resulted in a strong accumulation of the Non-Stop ProtA (Fig 5A). In contrast, *SKA1* overexpression had no effect on the degradation of XUTs (Fig 5B), yet was able to complement the *SKA1* deletion, showing that it did not impair the function of the SKI complex for the degradation of ribosome-free RNA sequences. These results are fully consistent with a model where Ska1 antagonizes the formation of ribosome-SKI complexes, which are involved in the NSD pathway as well as the degradation of translated RNA sequences (Schmidt *et al*, 2016). Therefore, we postulated that *SKA1* overexpression should promote the formation of the small complex visualized in a sucrose gradient to the detriment of the SKI/80S particle. To test this hypothesis, we examined the sedimentation of Ski3-TAP in a sucrose gradient when *SKA1* is overexpressed. In agreement with our hypothesis, we observed that the Ski3-TAP sedimentation profile shifts from the 80S to the region were Ska1-TAP normally sediments (Fig 5C), *i.e.* lighter fractions around the 40S peak (see Fig 1D).

**Figure 5.**
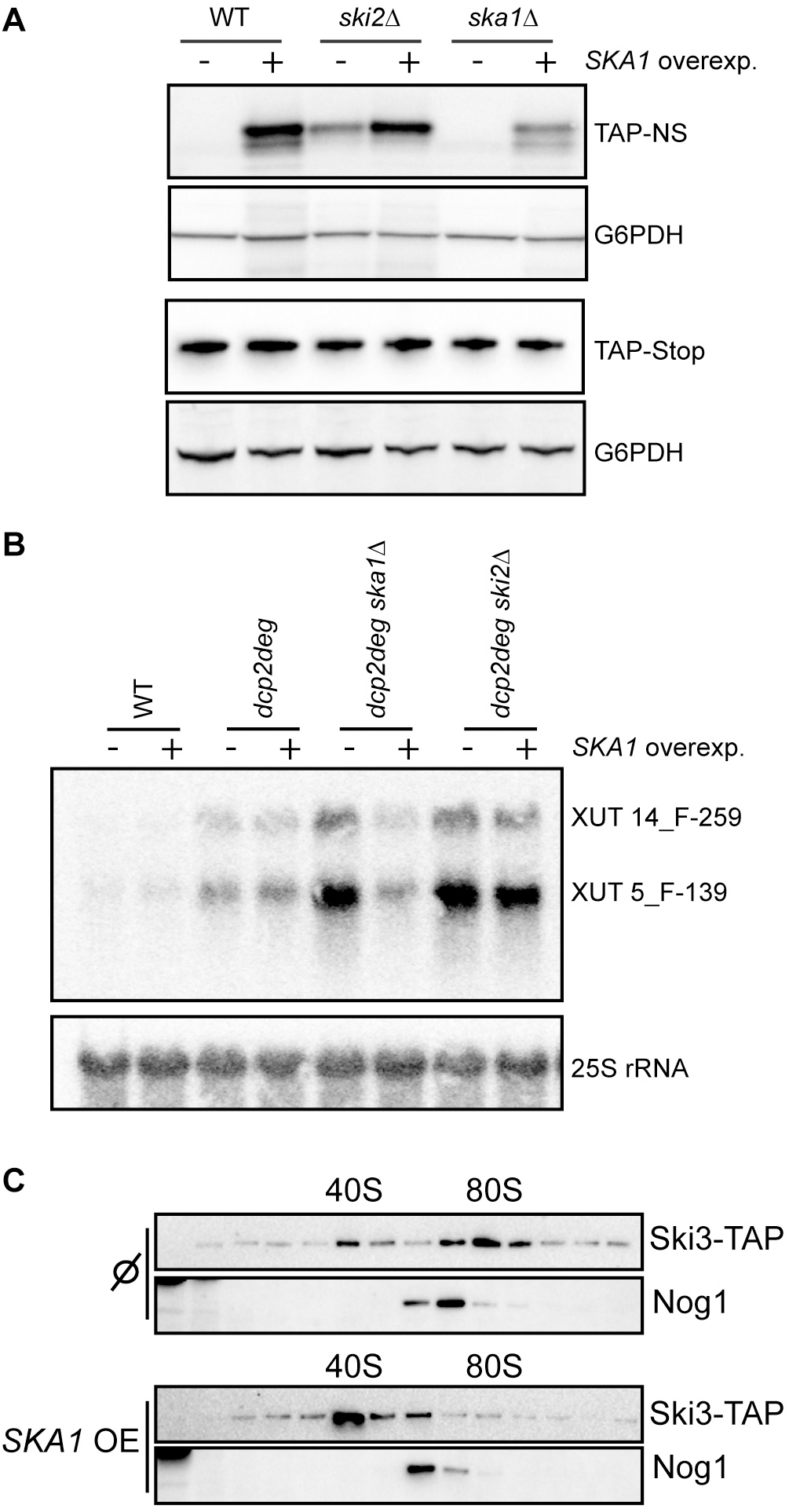
Ska1 overexpression affects Non-Stop mRNA decay. **A.** Total cellular extracts from the strains transformed with the TAP-Non-Stop or the TAP-Stop reporter genes as control were separated on a 10% SDS-PAGE polyacrylamide gel. Protein A was revealed by Western blot using a PAP antibody. A loading control was done using anti-G6PDH antibody. **B.** Northern blot with total RNAs from strains transformed with the empty vector pCM190 or the pCM190-SKA1 plasmid (“*SKA1* overexp.”) and treated with auxin for 2h, hybridized with probes for XUTs 14_F-259 and 5_F-139. A 25S-specific [^32^P]-radiolabeled oligonucleotide was used for a loading control. **C.** Total cellular extracts from cells expressing Ski3-TAP and transformed with the empty pCM190 vector (Ø) or with pCM190-SKA1 vector (*SKA1* OE) were separated on a sucrose gradient (10-50%) by ultracentrifugation. Each fraction was analyzed by Western blot using PAP for the detection of the TAP fusion proteins, as indicated. An anti-Nog1 antibody was used to mark the 60S fractions.

## Discussion

### The SKI-Ska1 complex is involved in the 3’-5’ degradation of mRNA 3’UTRs

It has been known for a long time that the SKI complex operates in the cytoplasm as an exosome co-factor to degrade RNAs from their 3’ ends. The SKI complex is not only involved in the general mRNA degradation pathway but also in aberrant mRNA degradation pathways such as the NSD or the NGD (Anderson & Parker, 1998; van Hoof *et al*, 2002). The recent finding that the SKI-exosome complex functions co-translationally and that the helicase activity of Ski2 is activated by its direct association with the ribosome (Schmidt *et al*, 2016) created a paradox because it is at odds with previous observations showing that the SKI complex is required for the efficient 3’-5’ degradation of mRNA 3’UTRs, mostly devoid of ribosomes (Anderson & Parker, 1998). The context in which the SKI complex operates for this specific function thus required further characterization. Using affinity purification followed by quantitative mass spectrometry analyses, we identified a new component, Ska1, specifically associated with a subpopulation of the SKI complex (Fig 1; Fig EV2). We demonstrated that this factor assists the SKI complex in the general mRNA degradation pathway but is not involved in the Non-Stop mRNA decay mechanism (Fig 2). The synergistic growth defect observed upon deletion of *SKA1* and *DCP2* was consistent with a role of Ska1 in the general mRNA decay (Fig 2D). We also showed that Ska1 might associate with ribosomes only indirectly by the intermediate of mRNAs (Fig 1). Furthermore, we found that, when the 5’-3’ degradation pathway was impaired by a depletion of Dcp2, Ska1 and the SKI complex were required for the efficient degradation of RNA regions devoid of ribosomes, such as 3’UTRs (Fig 2C) or cytoplasmic non-coding RNAs (Fig 3I, 4E and 4F). In contrast to Ski2, Ska1 is not required for the efficient degradation of short 3’UTRs (Fig 3B, 3D and S3B) or non-stop substrates, such as mRNAs prematurely cleaved and polyadenylated within their open reading frames (Fig 4A to 4D). Finally, we showed that overexpression of *SKA1* competes with the interaction between the SKI complex and the ribosome (Fig 5). We propose a model in which the general 3’-5’ mRNA degradation pathway could act in two steps after deadenylation (Fig 6). First, a Ska1-associated SKI complex would assist the exosome to degrade ribosome-free RNAs, such as 3’UTR mRNA regions, until it reaches the coding regions where it would encounter ribosomes. At this point, there are at least two models: the Ska1-SKI-exosome is exchanged for a “Ska1-free” SKI-exosome complex, which can directly interact with the ribosome; alternatively, Ska1 dissociates from the SKI-exosome complex, allowing it to directly interact with the ribosome while remaining on the RNA being degraded (Fig 6).

**Figure 6.**
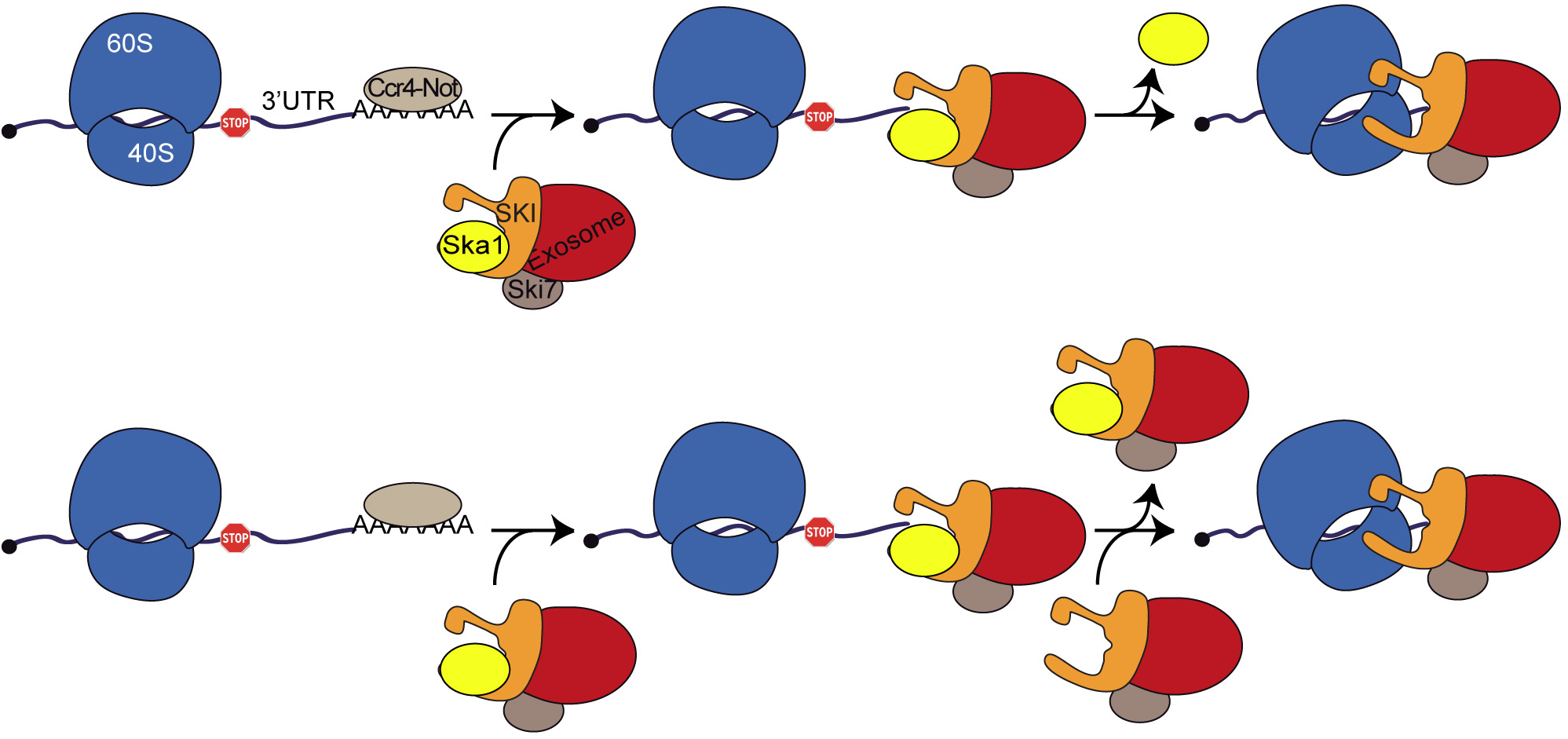
Model for SKI-ribosome and ribosome-independent SKI complexes functions. In a first step, the Ska1 associated SKI complex degrades the 3’UTR independently of the ribosome; in a second step, when the SKI/Ska1 complex reaches the coding sequence, there is an exchange between Ska1 and the ribosome for its association with the SKI complex (top). Alternatively (bottom), the Ska1-SKI complex is exchanged for the simple SKI complex, which can directly interact with the last ribosome on the mRNA.

The question arises as to know at what point the switch from the Ska1-SKI complex to the ribosome-SKI complex occurs. In many instances, (see Fig 3B and S3B), we observed that Ska1 was not required for the activity of the SKI complex on mRNAs with 3’UTRs shorter than about 60 nucleotides. This was confirmed by a genome-wide analysis, which showed that Ska1 had little effect on the 3’-5’ degradation of this class of mRNAs (Fig 3D and see below). Figure S4B shows examples of mRNAs with intermediate 3’UTR lengths (between ∼70 to 100 nucleotides). In all cases, similar profiles of 3’-degradation intermediates were observed between the *dcp2deg ski2*Δ and *dcp2deg ska1*Δ mutants until a distance of 55 to 67 nucleotides from the first nucleotide after the stop (sizes measured from the first nucleotide after the stop codon), from which Ska1 was no longer required (no accumulation of degradation intermediates) while Ski2 was. Interestingly, if one estimates the length of the RNA expected to be protected by the ribosome-SKI-exosome complex, one finds similar figures. More specifically, the ribosome on the termination codon has been shown to protect about 12 nucleotides following the stop codon (18 nucleotides when counting from the first nucleotide in the P site) (Archer *et al*, 2016). Adding the length of the RNA fragment protected *in vitro* by the SKI complex (9-10 nucleotides) (Halbach *et al*, 2013)) and by the exosome (31-33 nucleotides) (Malet *et al*, 2010) gives an expected range of protection between 52 and 55 nucleotides, counting from the first nucleotide following the stop codon. This length range, when compared to the 55-67 nucleotides distance from the stop codon from which Ska1 becomes dispensable (Fig EV4B) suggests that Ska1 is no longer required as soon as the SKI complex, which progresses from 3’-5’ along with the processive exosome (Dziembowski *et al*, 2007), comes sufficiently close to the last ribosome sitting on the stop codon to directly interact with it.

### What is the precise molecular function of Ska1?

We have shown in this work that Ska1 is required for the activity of the SKI-exosome complex during the degradation of ribosome-free RNA sequences. However, the precise molecular function of Ska1 within the Ska1-SKI-exosome complex remains unknown. Ska1 is a highly basic (pI: 9.65) protein of 32.3 kDa. Although homologs can be found in other fungi (SGD projet; https://www.yeastgenome.org/cache/fungi/YKL023W.html), it does not harbor any known protein domain and is therefore unlikely to have an enzymatic activity.

The SKI complex has been shown to have a strong affinity for ribosomes carrying 3’ RNA overhangs that are likely to be abundant in cells and thus susceptible to sequester most SKI complexes (Schmidt et al., 2016). We have shown that the overexpression of *SKA1* is able to divert the bulk of SKI complexes from interacting directly with ribosomes (Fig 5). It is possible that the major role of Ska1 is to prevent a fraction of the SKI complex from interacting with ribosomes, making it available for assisting the degradation of ribosome-free RNA sequences. According to this hypothesis, the only function of Ska1 would be to act as an antagonist of the SKI-ribosome interaction. Yet, the determination of the structure of the SKI-ribosome complex showed that the interaction between Ski2 and the 40S is important to activate the Ski2 helicase (Schmidt *et al*, 2016). This raises the possibility that Ska1 would play a similar role when it interacts with the SKI complex. Further experiments will be required to verify this possibility.

## Materials and Methods

### Yeast strains and plasmids

Yeast strains and plasmids used in this study are listed in Appendix Table S1 and S2, respectively. The yeast strains generated for this study were obtained by transformation of genomic PCR products. The details for each construction are available upon request. The *URA3* marker present in TAP-NS and TAP-Stop plasmids was replaced by a *LEU2* marker using the following strategy. The *LEU2* gene from pRS305 was amplified by PCR using oligonucleotides QD139 and QD140 and transformed in yeast cells with plasmid TAP-NS or TAP-Stop linearized with EcoRV for recombination. pCM190-SKA1: the *SKA1* gene was amplified by PCR using oligonucleotides EZ091 and EZ092 from BY4741 genomic DNA and the fragment cut with BamHI and NotI and subcloned into pCM190 plasmid cut with the same enzymes.

### Polysome gradients, proteins extraction and western blotting

Polysome extracts and sucrose gradients were performed as described (Defenouillère *et al*, 2013). Total proteins extracts were obtained by alkaline treatment using 5 A_600_ mid-log phase yeast cells. The extracts were separated on 10% SDS-PAGE polyacrylamide gel. The proteins were transferred on nitrocellulose membrane using a Biorad semi-dry machine. The membrane was hybridized with appropriate antibody dilutions as mentioned in Appendix Table 3.

### RNA extraction and Northern blot analysis

Total RNA was purified from 12 A_600_ mid-log phase yeast cells using the hot acid phenol protocol (Collart & Oliviero, 2001). RNAs were separated on 1% agarose gel, transferred on nylon membranes (Hybond N+, Amersham) and probed with ^32^P-labeled oligonucleotides or with ^32^P-labeled RNA probe synthetized according to the Ambion MAXIscript^®^ Kit procedure using the oligonucleotides listed in Appendix Table 4.

### Libraries construction

Three independent biological replicates were generated for wild-type, single and double mutant strains. Cells were grown to exponential phase and treated with auxin (final concentation 0.1mM) for 2h. Reference aliquots of *Schizosaccharomyces pombe* were added and total RNA isolated. Five µg of total RNA were treated with Turbo DNAseI for 20 min. at 37°C. After phenol/chloroform extraction, RNAs were precipitated with ethanol and resuspended in water. A depletion of the ribosomal RNA was performed using Ribo-Zero Gold rRNA removal kit for yeasts (Illumina). Removal of the rRNA was verified using a Bioanalyzer. An *in vitro* polyadenylation was performed with 4 units of PAP in the presence of 50 µM cordycepin and 1mM ATP. After phenol/chloroform extraction, RNAs were precipitated with ethanol and resuspended in water. The 3’mRNA-Seq libraries were constructed using the Lexogen QuantSeq 3’ mRNA-Seq Library Prep Kit for Illumina (Rev) with Custom Sequencing Primer according to manufacter’s instruction. The RNASeq libraries were made using 5 µg of total RNA treated with Turbo DNAse and Ribo-Zero as described above. The libraries were constructed using TruSeq stranded mRNA LT sample prep Kits (Illumina) according to the manufacturer’s instruction. The total read counts for each library are indicated in Appendix Table 5. The reproducibility of the experiments was tested by Principal Component Analysis (PCA) for three independent biological replicates per strain. The replicates for each mutant strain clustered together and aside from the other strains (Fig EV5A). The genome-wide analysis for each replicate before merging is shown in Fig EV5B.

### **Bioinformatics analyses** are described in the Appendix

The sequencing data are available from European Nucleotide Archive (accession number PRJEB27292, http://www.ebi.ac.uk/ena/data/view/PRJEB27292) for all TTS sequencing samples under the accession codes ERS2548668 to ERS2548690 and for all RNA sequencing samples under the accession codes ERS2548828 to ERS2548845.

### Affinity purifications coupled to mass spectrometry

Affinity purifications were performed using 4 liters of culture at 2 A_600_ as described in (Defenouillère *et al*, 2013). Briefly, proteins were digested in solution and then peptides were analyzed using an LTQ-Orbitrap Velos. Protein identification and comparative label-free quantification were performed using the MaxQuant suite (version 1.5.5.1) and the Andromeda search engine. Quantifications were done using the algorithm integrated into MaxQuant to calculate Label Free Quantification intensities (Appendix).

## Acknowledgments

We thank colleagues mentioned in the text for their gifts of material. We are grateful to Julia Chamot-Rooke and Mariette Matondo from the proteomic platform of the Pasteur Institute for the availability of the Orbitrap instrument and to Odile Sismeiro and Jean-Yves Coppée from the Biomics pole of the Pasteur Institute for running the sequencing libraries on the Illumina machine. We thank the members of the lab, notably Cosmin Saveanu and Gwenael Breard for reading the manuscript and particularly Maïté Gomard who recently joined the team. E. Z and E.D were supported by ANR-14-CE-10-0014-01 and ANR-17-CE11-0049-01 grants from the Agence Nationale de la Recherche (ANR) respectively. This work was supported by ANR-14-CE-10-0014-01 and ANR-17-CE11-0049-01 grants, the Pasteur Institute and the Centre National de la Recherche Scientifique. B.T. was supported by a grant from the Natural Sciences and Engineering Research Council of Canada.

## Author contributions

E.Z., A.D., B.T., E.D., A.J. and M.F-R. conducted the experiments, A.N. generated and analyzed the mass spectrometry data. V.K. analyzed the RNA Seq data. M.F-R. and A.J. wrote the paper and designed the experiments. All authors reviewed the results and approved the final version of the manuscript.

## Expanded View Figure Legends

**Figure EV1.**
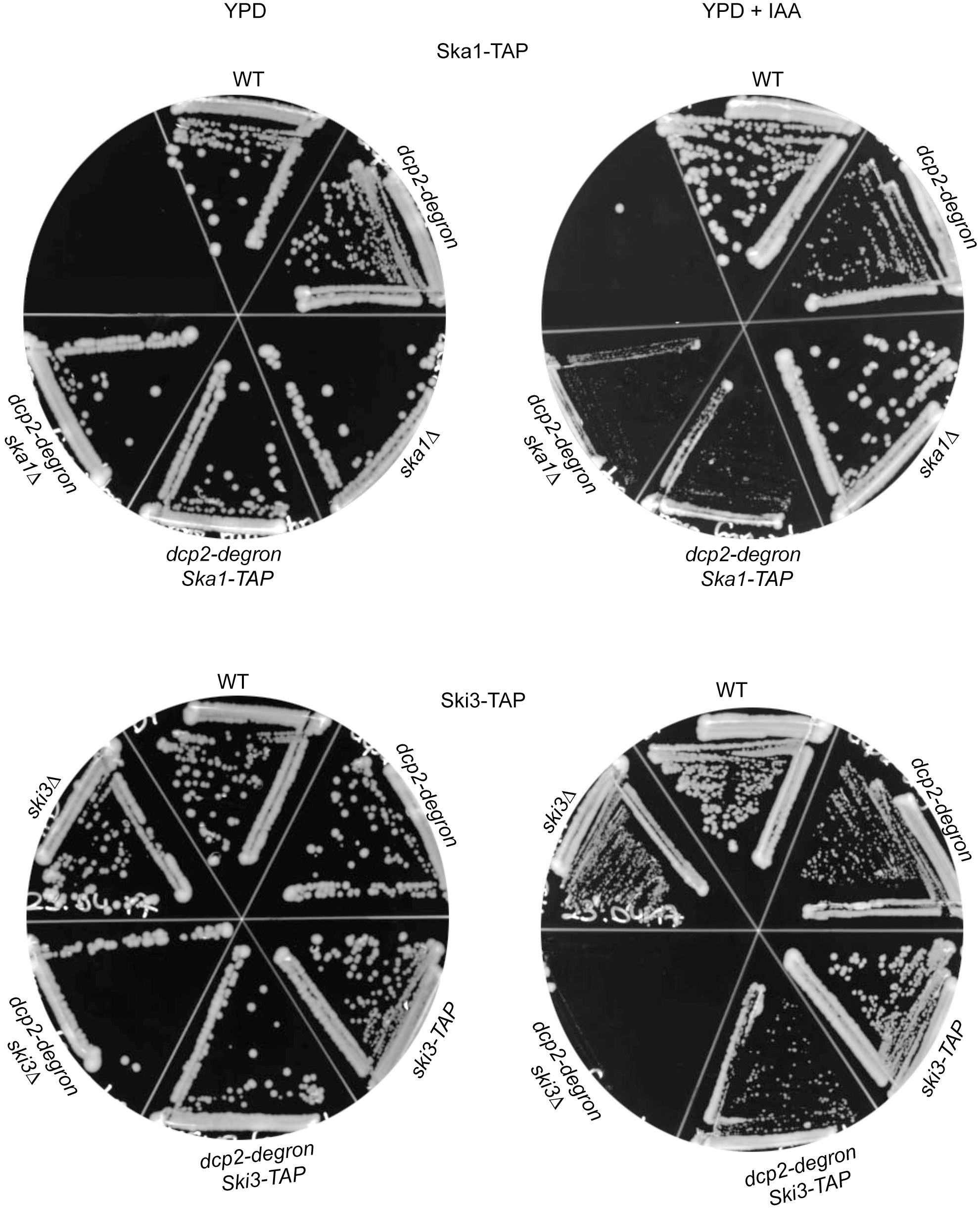
Ski3-TAP is functional and Ska1-TAP is partially functional. Single (*ski3Δ, ska1Δ,* Ska1-TAP, Ski3-TAP, *dcp2deg*) or double (*dcp2deg ski3Δ, dcp2deg ska1Δ, dcp2deg* Ski3-TAP, *dcp2deg* Ska1-TAP) mutant strains were spread on rich medium plates with or without auxin (IAA; 0.1 mM) and incubated for 48h at 30°C.

**Figure EV2.**
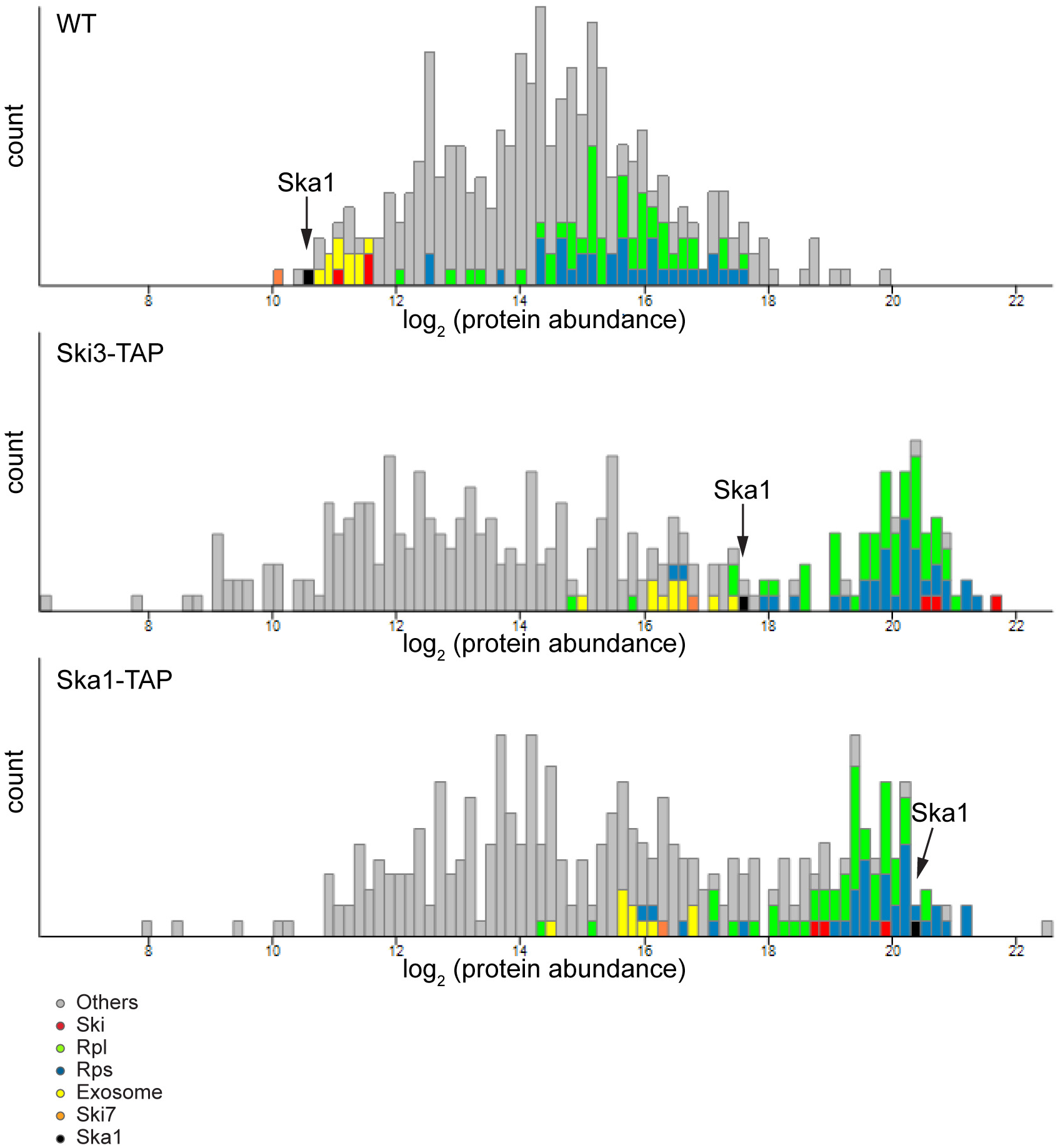
Relative protein abundance. Each stacked histogram shows the median of the label free quantification (count; arbitrary units) of each protein in log2 scale in purified SKI3-TAP and purified SKA1-TAP as well as in the total extract of WT cells (data from Nagaraj et al, 2012). Purified samples (Ski3-TAP, Ska1-TAP) showed enriched proteins (colors, as indicated) when compared to the WT total extracts.

**Figure EV3.**
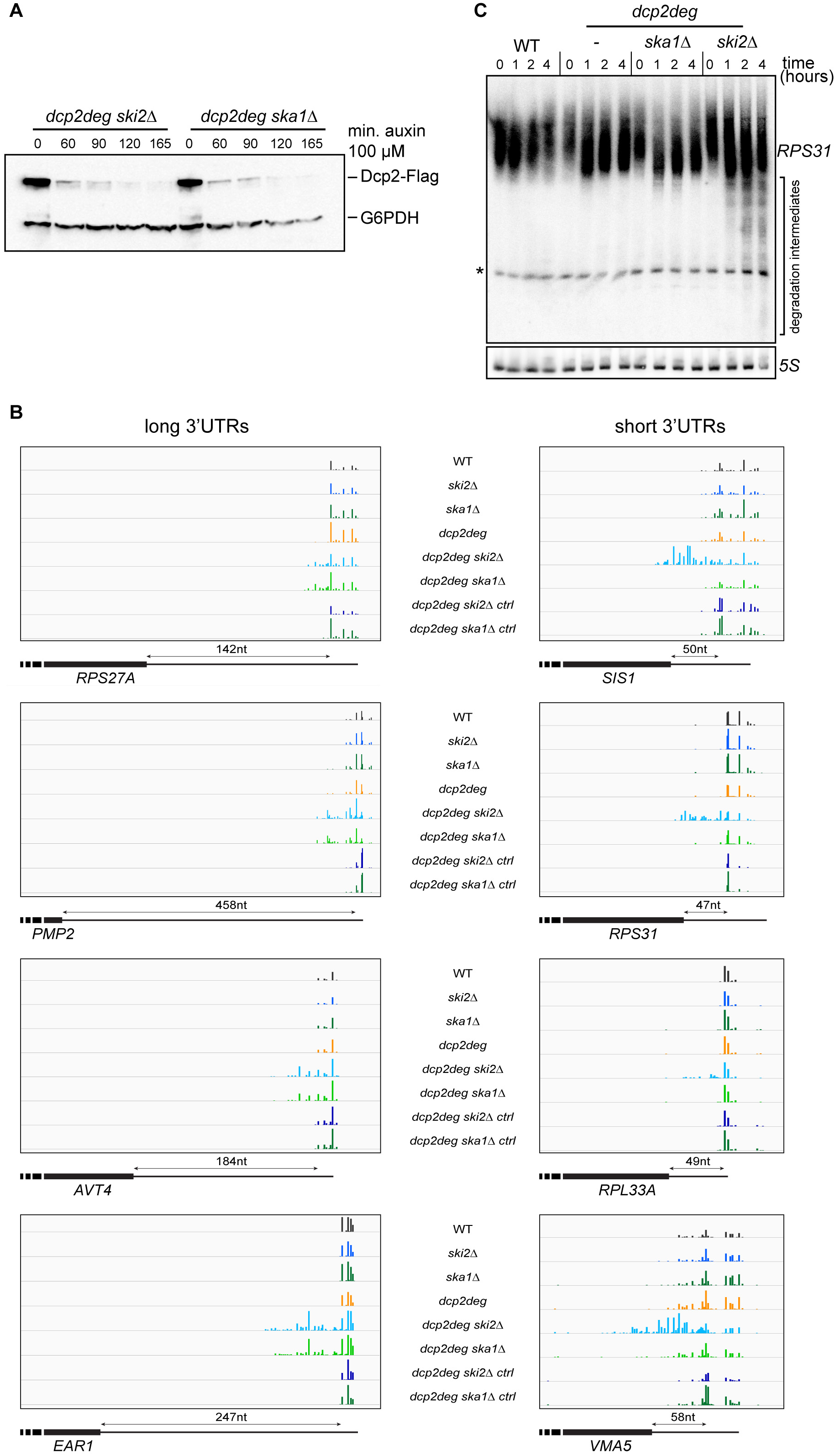
Examples of RNA intermediates accumulated in mutant strains. **A.** Total cellular extracts from the *dcp2deg ski2*Δ and *dcp2deg ska1*Δ strains treated with auxin for different times were separated on 10% SDS-PAGE polyacrylamide gel. The Dcp2 protein fused to Flag tag was revealed by Western blot anti-FLAG antibody. A loading control of the total protein amounts was done using anti-G6PDH antibody. **B.** Visualization (using IGV) of the 3’ends, with or without (ctrl lanes) the in vitro polyadenylation step, for long (left panel) or short 3’UTR (right panel) containing gene loci. **C.** The extent of the 3’-end trimming of the RPS31 mRNA in the *dcp2deg ski2*Δ double mutant depends on the length of auxin treatment WT, *dcp2deg, dcp2deg ski2*Δ or *dcp2deg ska1*Δ cells were exposed to auxin (0,1 mM) for 0, 1, 2 or 4 hours, as indicated. Total RNAs were extracted and treated with RNase H in the presence of an oligonucleotide (EsD11) complementary to a sequence located close to the end of the *RPS31* ORF (from -75 to -58 nucleotides from the stop codon). RNA was separated on a 6% polyacrylamide gel, transferred to a nylon membrane and hybridized with a [^32^P]-radiolabeled RNA probe complementary to sequences -57 to -29 relative to the stop codon (RNA was generated by T7 *in vitro* transcription in the presence of [^32^P]-UTP and the hybridized oligonucleotides EsD12 and AJ529). The asterisk marks a crosshybridizing RNA migrating in the tRNA region of the gel. Note that the length of the main *RPS31* species (noted *RPS31* on the gel) shortens upon auxin treatment, denoting accumulation of deadenylated *RPS31* after Dcp2 depletion. The lengths of the smears below the main *RPS31* species (labeled “degradation intermediates”), only visible in the *dcp2deg ski2*Δ double mutant, as expected for 3’ trimmed species accumulating in this “short 3’UTR” gene (see Figure EV3B), gets longer with the length of auxin depletion.

**Figure EV4.**
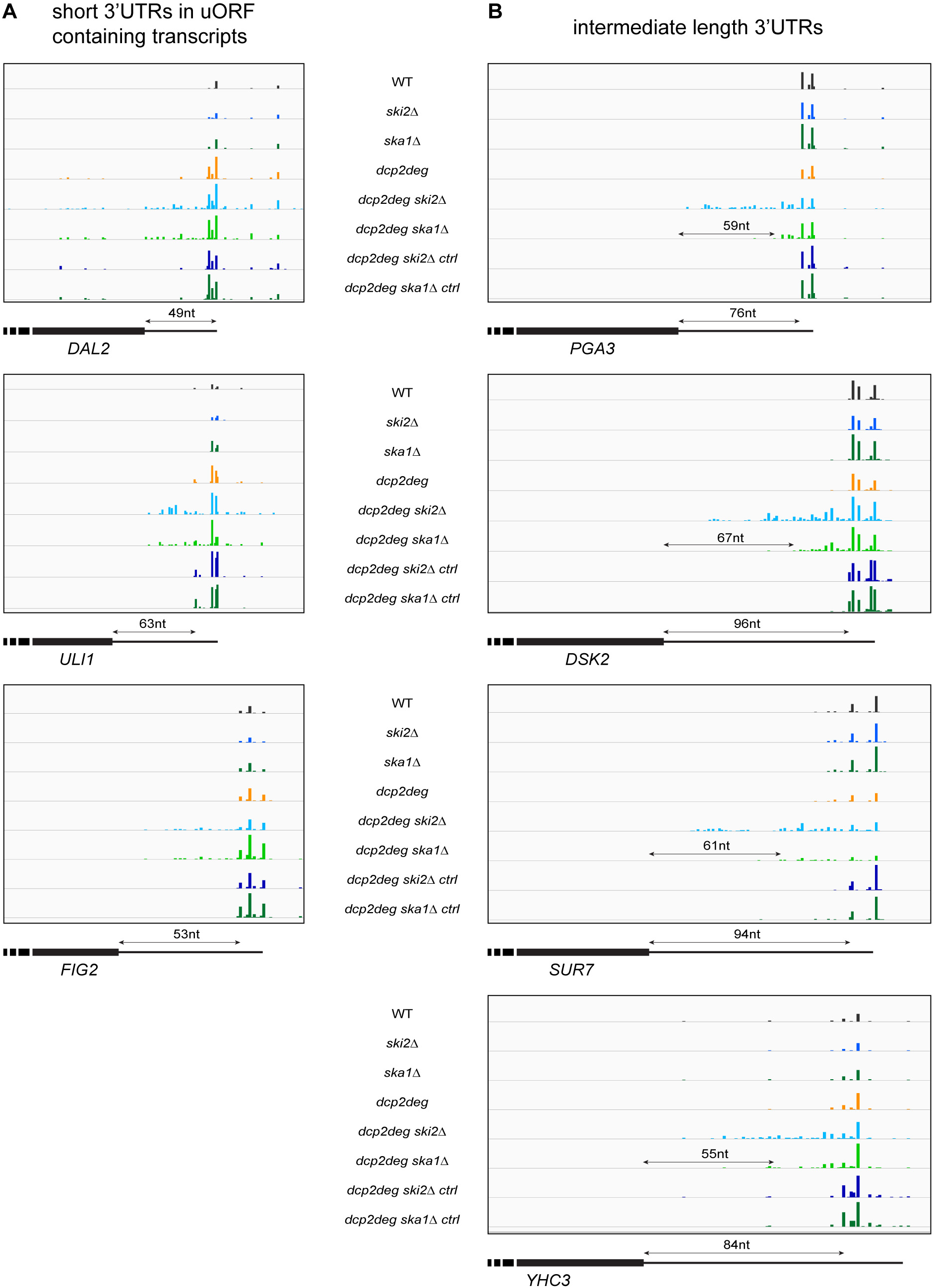
Visualization of 3’ degradation intermediates. Same as Figure 3 A-B for genes with short 3’UTR but containing uORFs (**A)** or genes with intermediates length 3’UTR (**B)** The double arrowed lines show sequence length in nucleotides (nt).

**Figure EV5.**
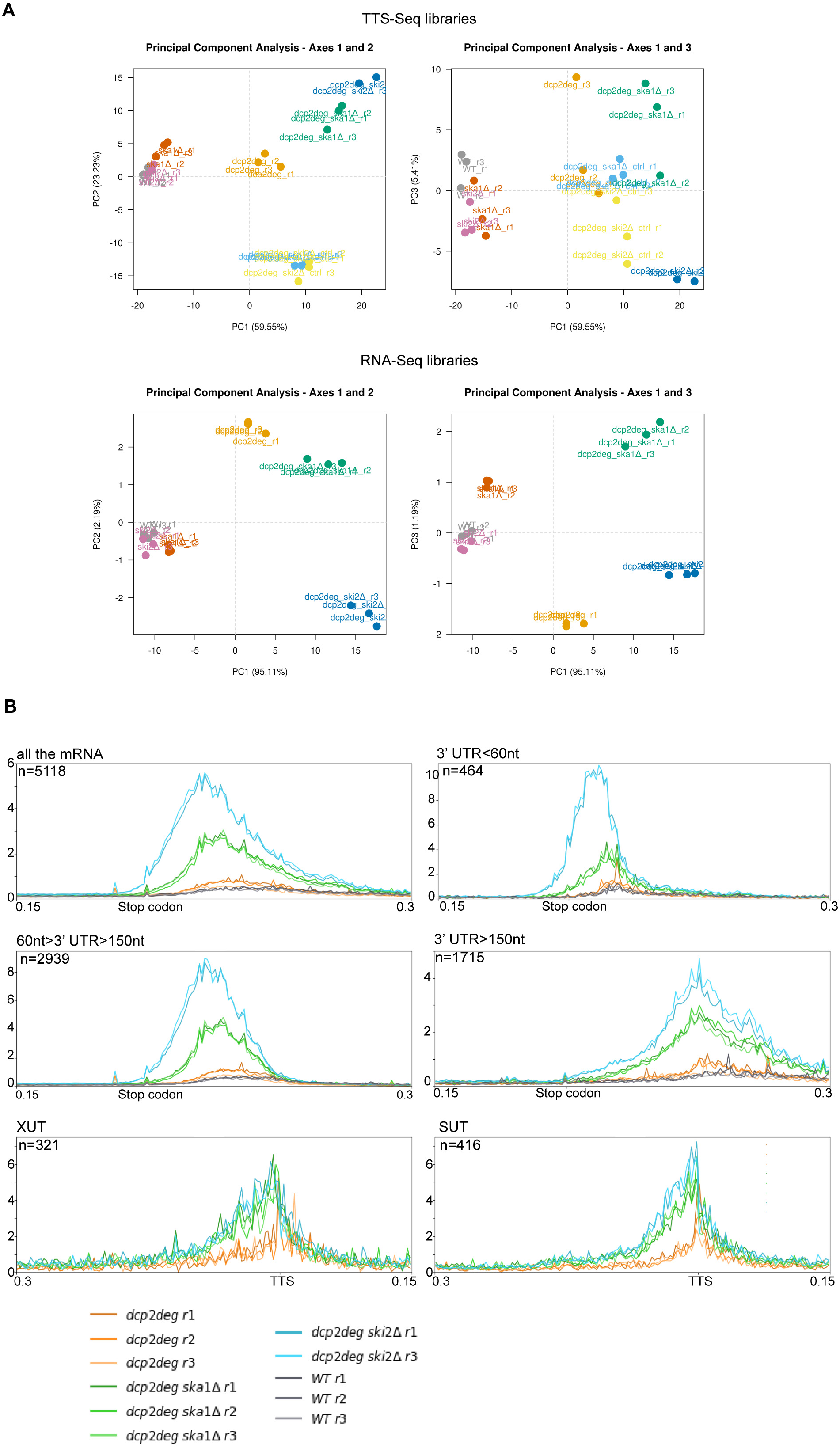
**A.** Dimensionality reduction in TTS-seq and RNA-seq libraries analyses generated by SARTools package (see Appendix). The plots show the first three dimensions of PCA for all conditions tested. In left: PC1 vs PC2. In right: PC1 vs PC3. **B.** Visualization of the genome-wide analyses, as in Figure 3, but for all individual replicates.

